# ERGA-BGE genome of *Coenonympha oedippus*: an IUCN endangered European butterfly species occurring in two ecotypes

**DOI:** 10.1101/2025.02.10.637396

**Authors:** Tatjana Čelik, Tjaša Lokovšek, Elena Bužan, Astrid Böhne, Rita Monteiro, Rosa Fernández, Nuria Escudero, Marta Gut, Laura Aguilera, Francisco Câmara Ferreira, Fernando Cruz, Jèssica Gómez-Garrido, Tyler S. Alioto, Leanne Haggerty, Fergal Martin, Chiara Bortoluzzi

## Abstract

The reference genome of the False ringlet (*Coenonympha oedippus*) will serve as a valuable resource for uncovering the genetic mechanism underlying the species′ adaptability to two ecologically distinct habitats. Through this genome we might be able to determine whether (i) each ecotype is monophyletic, indicating that the ecological divergence represents an early stage of speciation, (ii) the ecotypes have evolved through divergent evolution of habitat preference, or (iii) the differences between ecotypes are solely due to phenotypic plasticity or epigenetic variation. This reference genome is also a prerequisite for the planning, design, and implementation of conservation measures for this endangered species, taking into account its intraspecific diversity. Furthermore, it holds broader implications for population genomic studies of the species-rich genus *Coenonympha*, which includes some of the most endangered butterfly taxa in Europe. The complete genome sequence was assembled into 30 contiguous chromosomal pseudomolecules (sex chromosomes included). This chromosome-level assembly encompasses 0.39 Gb, composed of 385 contigs and 62 scaffolds, with contig and scaffold N50 values of 2.8 Mb and 14.2 Mb, respectively.

## Introduction

*Coenonympha oedippus*, also known as False ringlet or Barjanski okarček in Slovenian, is a Eurasian butterfly species from the subfamily *Satyrinae*. The species shows remarkable ecotype differences in its European distribution area, which are reflected in the different larval host plants (Čelik et al., 2015), wing morphology (Jugovic et al., 2018), and neutral genetic differentiation (Zupan et al., 2021). Given the phylogeographic distribution of the species in Europe (Després et al., 2019), adaptive genetic differentiation is expected. The recent evolutionary history of *Coenonympha* butterflies has shown that *C. oedippus* (together with the genus *Triphysa*) belongs to an early divergent clade that split from other *Coenonympha* species about 13 million years ago (Greenwood et al., 2025).

*Coenonympha oedippus* was last assessed for the IUCN Red List of Threatened Species in 2009, during which it was categorised as Endangered under criterion A2c according to Europe’s assessment criteria (van Swaay, 2010). The species is also listed in Annexes II and IV of the EU Habitats Directive (Council Directive 92/43/EEC) and its conservation status is assessed as ‘Unfavourable’ (U2, U1) in most EU Member States (Article 17 reporting, 2013–2018). The initial conservation translocations (i.e. reintroduction and reinforcements) of the species in Slovenia (2018–2022) highlighted the need to assess genetic diversity within the source and reintroduced populations to evaluate whether genetic variation needs to be increased through dedicated management strategies, such as genetic rescue or assisted gene flow (Čelik et al., 2024).

*Coenonympha oedippus* is an important indicator species that reflect the health status of oligotrophic wet grassland and fens. The larvae are nutritionally dependent on plant species that are dominant and characteristic of these grasslands and therefore could influence plant population dynamics (Čelik et al., 2021). They are also hosts for parasites (Dierks, 2006) and are an important food source for many predatory invertebrates, birds, and small mammals in addition to the other three life stages (egg, pupa, butterfly) (Lhonoré, 1998). The complex life cycle of *C. oedippus*, with four developmental stages, each with specific habitat requirements at different scales - from microhabitat to landscape - places the species at the centre of restoration and management planning, which also benefits wider biodiversity (Čelik et al., 2015, 2024).

The generation of this reference resource was coordinated by the European Reference Genome Atlas (ERGA) initiative’s Biodiversity Genomics Europe (BGE) project, supporting ERGA’s aims of promoting transnational cooperation to promote advances in the application of genomics technologies to protect and restore biodiversity (Mazzoni et al., 2023).

## Materials & Methods

ERGA’s sequencing strategy includes Oxford Nanopore Technology (ONT) and/or Pacific Biosciences (PacBio) for long-read sequencing, along with Hi-C sequencing for chromosomal architecture, Illumina Paired-End (PE) for polishing (i.e. recommended for ONT-only assemblies), and RNA sequencing for transcriptome profiling, to facilitate genome assembly and annotation.

### Sample and Sampling Information

On 15 June 2023, Tatjana Čelik and Tjaša Lokovšek sampled 5 female specimens of *Coenonympha oedippus*. The specimens were identified by Tatjana Čelik based on morphology without an identification key, as the expert knows the species well, having researched it for 20 years. The 5 specimens were collected in Babiči-Zupančiči, Slovenska Istra (Slovenia). Sampling was performed under permission 35601-140/2009 – 6 issued by the Ministry of the Environment and Spatial Planning of the Republic of Slovenia. Specimens were hand-collected and then euthanized using a light mechanical compression of the thorax. At the collection site, each specimen was snap-frozen and subsequently preserved at -80°C until DNA extraction.

### Vouchering information

Physical reference materials from a proxy female individual (i.e., an individual different from the one used for sequencing) have been deposited in the Prirodoslovni muzej Slovenije https://www.pms-lj.si/, Ljubljana (Slovenia) under proxy voucher ID PMSL-Invertebrata-22.

Frozen reference tissue material of a whole organism (female) is available in the Prirodoslovni muzej Slovenije https://www.pms-lj.si/, Ljubljana (Slovenia) under proxy tissue voucher ID PMS_TIS_14.

### Data Availability

*Coenonympha oedippus* and the related genomic study were assigned to Tree of Life ID (ToLID) ‘ilCoeOedi1’ and all sample, sequence, and assembly information are available under the umbrella BioProject PRJEB77778. The sample information is available at the following BioSample accessions: SAMEA115177279 and SAMEA115177280. The genome assembly is accessible from ENA under accession number GCA_964304575.1 and the annotated genome is available through the Ensembl Beta website (https://projects.ensembl.org/erga-bge/). Sequencing data produced as part of this project are available from ENA at the following accessions: ERX13168332, ERX13168333, and ERX12746377.

Documentation related to the genome assembly and curation can be found in the ERGA Assembly Report (EAR) document available at https://github.com/ERGA-consortium/EARs/tree/main/Assembly_Reports/Coenonympha_oedippus/ilCoeOedi1. Further details and data about the project are hosted on the ERGA portal at https://portal.erga-biodiversity.eu/data_portal/554486.

### Genetic Information

The estimated genome size, based on ancestral taxa, is 0.48 Gb. This is a diploid genome with a haploid number of 29 chromosomes (2n=58) and unknown sex chromosomes. All information for this species was retrieved from Genomes on a Tree (Challis et al., 2023).

### DNA/RNA processing

DNA was extracted from the whole organism using the Blood & Cell Culture DNA Midi Kit (Qiagen) following the manufacturer’s instructions. DNA quantification was performed using a Qubit dsDNA BR Assay Kit (Thermo Fisher Scientific), and DNA integrity was assessed using a Genomic DNA 165 Kb Kit (Agilent) on the Femto Pulse system (Agilent). The DNA was stored at +4°C until used.

RNA was extracted using an RNeasy Mini Kit (Qiagen) according to the manufacturer’s instructions. RNA was extracted from three different specimen parts: head, wing and abdomen. RNA quantification was performed using the Qubit RNA BR kit and RNA integrity was assessed using a Bioanalyzer 2100 system (Agilent) RNA 6000 Nano Kit (Agilent). The RNA was pooled equimolarly for the library preparation and stored at -80°C until used.

Library Preparation and Sequencing For long-read whole genome sequencing, a library was prepared using the SQK-LSK114 Kit (Oxford Nanopore Technologies, ONT), which was then sequenced on a PromethION 24 A Series instrument (ONT). A short-read whole genome sequencing library was prepared using the KAPA Hyper Prep Kit (Roche). A Hi-C library was prepared from thorax using the Dovetail Omni-C Kit (Cantata Bio), followed by the KAPA Hyper Prep Kit for Illumina sequencing (Roche). The RNA library was prepared from the pooled sample using the KAPA mRNA Hyper prep kit (Roche). The short-read libraries were sequenced on a NovaSeq 6000 instrument (Illumina).

In total 226x Oxford Nanopore, 75x Illumina WGS shotgun, and 183x HiC data were sequenced to generate the assembly.

### Genome Assembly Methods

The genome was assembled using the CNAG CLAWS pipeline (Gomez-Garrido, 2024). Briefly, reads were pre-processed for quality and length using Trim Galore v0.6.7 and Filtlong v0.2.1, and initial contigs were assembled using both NextDenovo v2.9.0, and Flye v.2.9.1 followed by polishing of the assembled contigs using HyPo v1.0.3, removal of retained haplotigs using purge-dups v1.2.6 and scaffolding with YaHS v1.2a. Finally, the Flye-assembled scaffolds were curated via manual inspection using Pretext v0.2.5 with the Rapid Curation Toolkit (https://gitlab.com/wtsi-grit/rapid-curation) to remove any false joins and incorporate any sequences not automatically scaffolded into their respective locations in the chromosomal pseudomolecules (or super-scaffolds). The contig corresponding to the complete W chromosome from the Nextdenovo assembly was used to replace the multiple scaffolds corresponding to the W chromosome in the Flye assembly. Finally, the mitochondrial genome was assembled as a single circular contig of 15,325 bp using the FOAM pipeline (https://github.com/cnag-aat/FOAM) and included in the released assembly (GCA_964304575.1). Summary analysis of the released assembly was performed using the ERGA-BGE Genome Report ASM Galaxy workflow (10.48546/workflow hub.workflow.1103.2).

### Genome Annotation Methods

A gene set was generated using the Ensembl Gene Annotation system (Aken et al., 2016), primarily by aligning publicly available short-read RNA-seq data from BioSample: SAMEA117648699 to the genome. Gaps in the annotation were filled via protein-to-genome alignments of a select set of clade-specific proteins from UniProt (Consortium, 2019), which had experimental evidence at the protein or transcript level. At each locus, data were aggregated and consolidated, prioritising models derived from RNA-seq data, resulting in a final set of gene models and associated non-redundant transcript sets. To distinguish true isoforms from fragments, the likelihood of each open reading frame (ORF) was evaluated against known metazoan proteins. Low-quality transcript models, such as those showing evidence of fragmented ORFs, were removed. In cases where RNA-seq data were fragmented or absent, homology data were prioritised, favouring longer transcripts with strong intron support from short-read data. The resulting gene models were classified into two categories: protein-coding, and long non-coding. Models that did not overlap protein-coding genes, and were constructed from transcriptomic data were considered potential lncRNAs. Potential lncRNAs were further filtered to remove single-exon loci due to their unreliability. Putative miRNAs were predicted by performing a BLAST search of miRBase (Kozomara et al., 2019) against the genome, followed by RNAfold analysis (Gruber et al., 2008). Other small non-coding loci were identified by scanning the genome with Rfam (Kalvari et al., 2018) and passing the results through Infernal (Nawrocki & Eddy, 2013). Summary analysis of the released annotation was carried out using the ERGA-BGE Genome Report ANNOT Galaxy workflow (10.48546/workflowhub.workflow.1096.1).

## Results

### Genome Assembly

The genome assembly has a total length of 397,595,348 bp in 62 scaffolds including the mitogenome (Figures 1 & 2), with a GC content of 37.3%. The assembly has a contig N50 of 2,773,664 bp and L50 of 45 and a scaffold N50 of 14,242,255 bp and L50 of 13. The assembly has a total of 323 gaps, totalling 65.7 kb in cumulative size. The single-copy gene content analysis using the Lepidoptera database with BUSCO (Manni et al., 2021) resulted in 98.2% completeness (97.5% single and 0.7% duplicated). 69.4% of reads k-mers were present in the assembly and the assembly has a base accuracy Quality Value (QV) of 50.4 as calculated by Merqury (Rhie et al., 2020).

**Figure 1.**
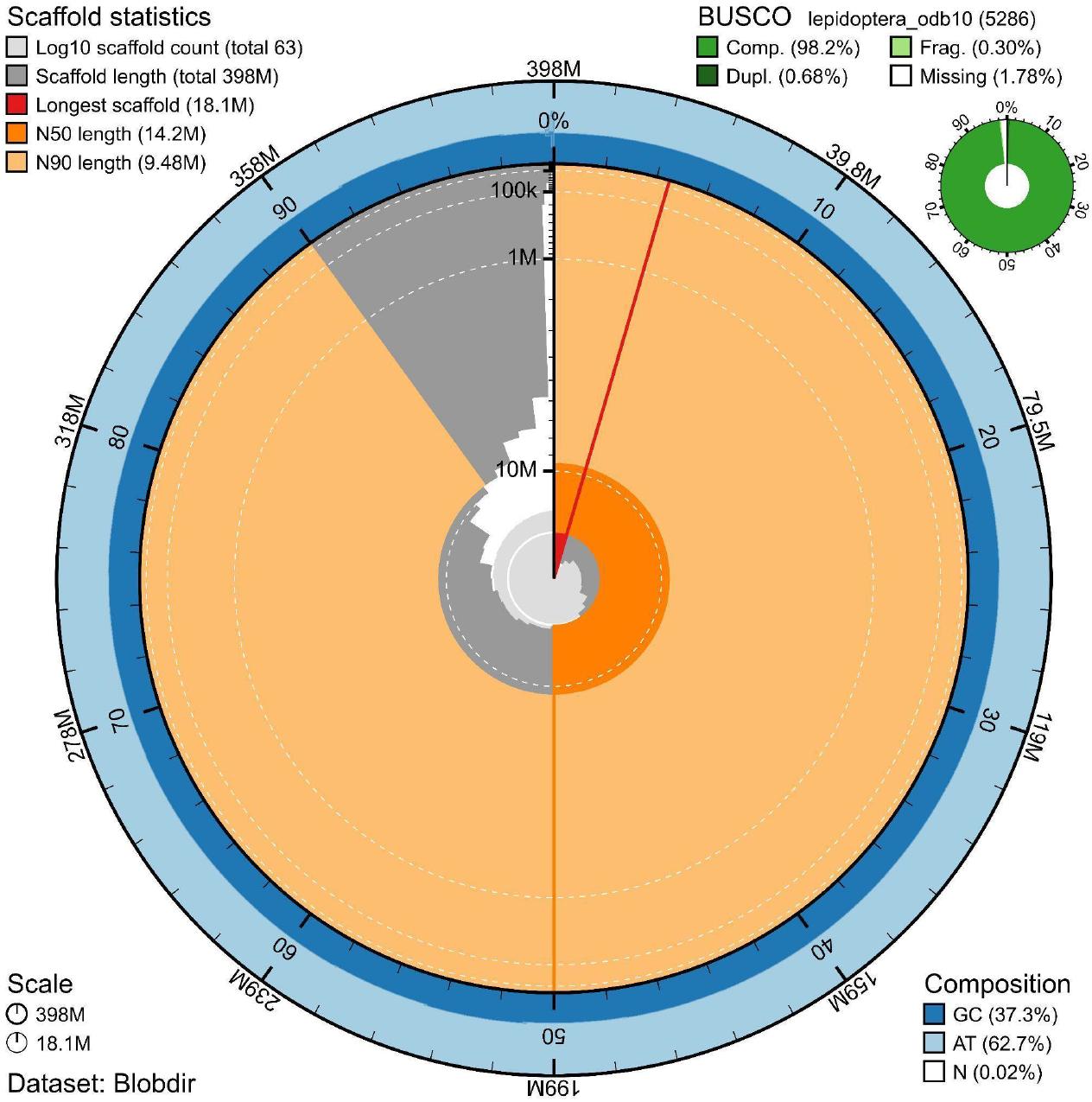
Snail plot summary of assembly statistics. The main plot is divided into 1,000 size-ordered bins around the circumference, with each bin representing 0.1% of the 397,595,348 bp assembly including the mitochondrial genome. The distribution of sequence lengths is shown in dark grey, with the plot radius scaled to the longest sequence present in the assembly (18.1 Mb, shown in red). Orange and pale-orange arcs show the scaffold N50 and N90 sequence lengths (14,242,255 and 9,479,849 bp), respectively. The pale grey spiral shows the cumulative sequence count on a log-scale, with white scale lines showing successive orders of magnitude. The blue and pale-blue area around the outside of the plot shows the distribution of GC, AT, and N percentages in the same bins as the inner plot. A summary of complete, fragmented, duplicated, and missing BUSCO genes found in the assembled genome from the Lepidoptera database (odb10) is shown in the top right.

**Figure 2.**
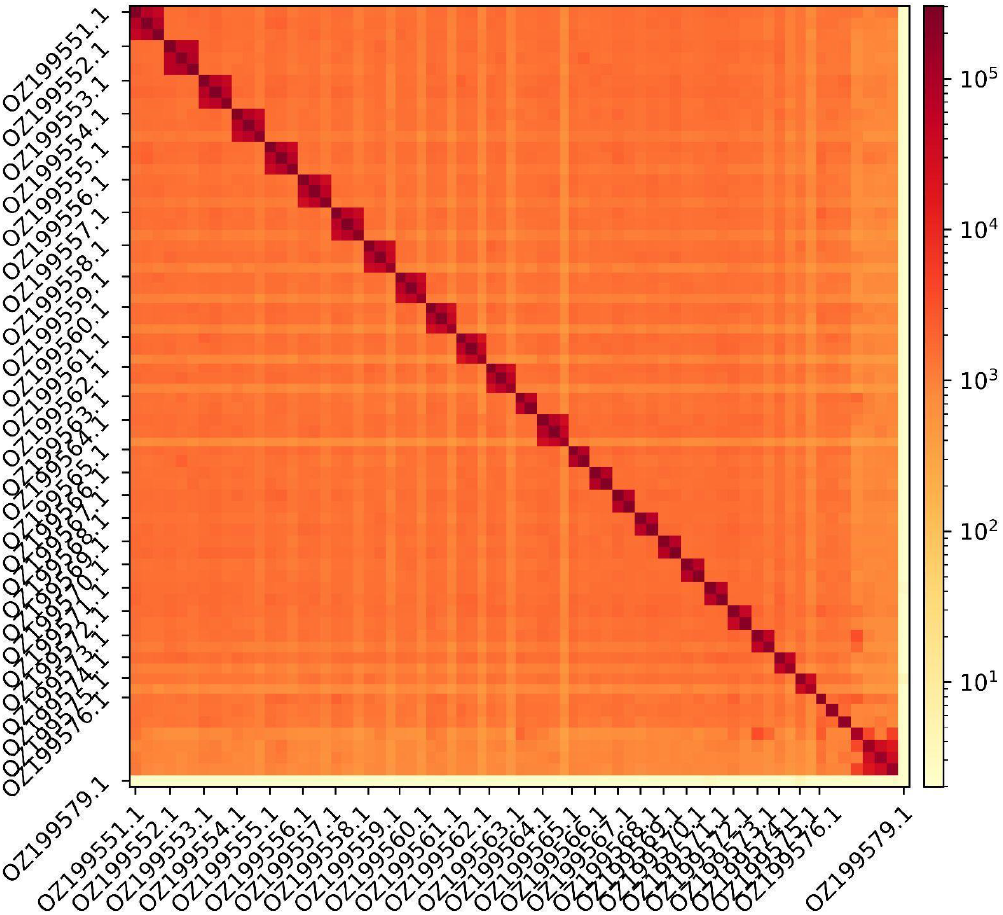
Hi-C contact map showing spatial interactions between regions of the genome. The diagonal corresponds to intra-chromosomal contacts, depicting chromosome boundaries. The frequency of contacts is shown on a logarithmic heatmap scale. Hi-C matrix bins were merged into a 100 kb bin size for plotting. Due to space constraints on the axes, only the GenBank names of the 26th largest chromosomes and the mitochondrial genome (GenBank name: OZ199579.1) are shown.

### Genome Annotation

The genome annotation consists of 12,720 protein-coding genes with associated 21,719 transcripts, in addition to 2,708 non-coding genes (Table 1). Using the longest isoform per transcript, the single-copy gene content analysis using the Lepidoptera odb10 database with BUSCO resulted in 93.2% completeness. Using the OMAmer Metazoa-v2.0.0.h5 database for OMArk (Nevers et al., 2025) resulted in 92.0% completeness and 89.9% consistency (Table 2).

**Table 1.**
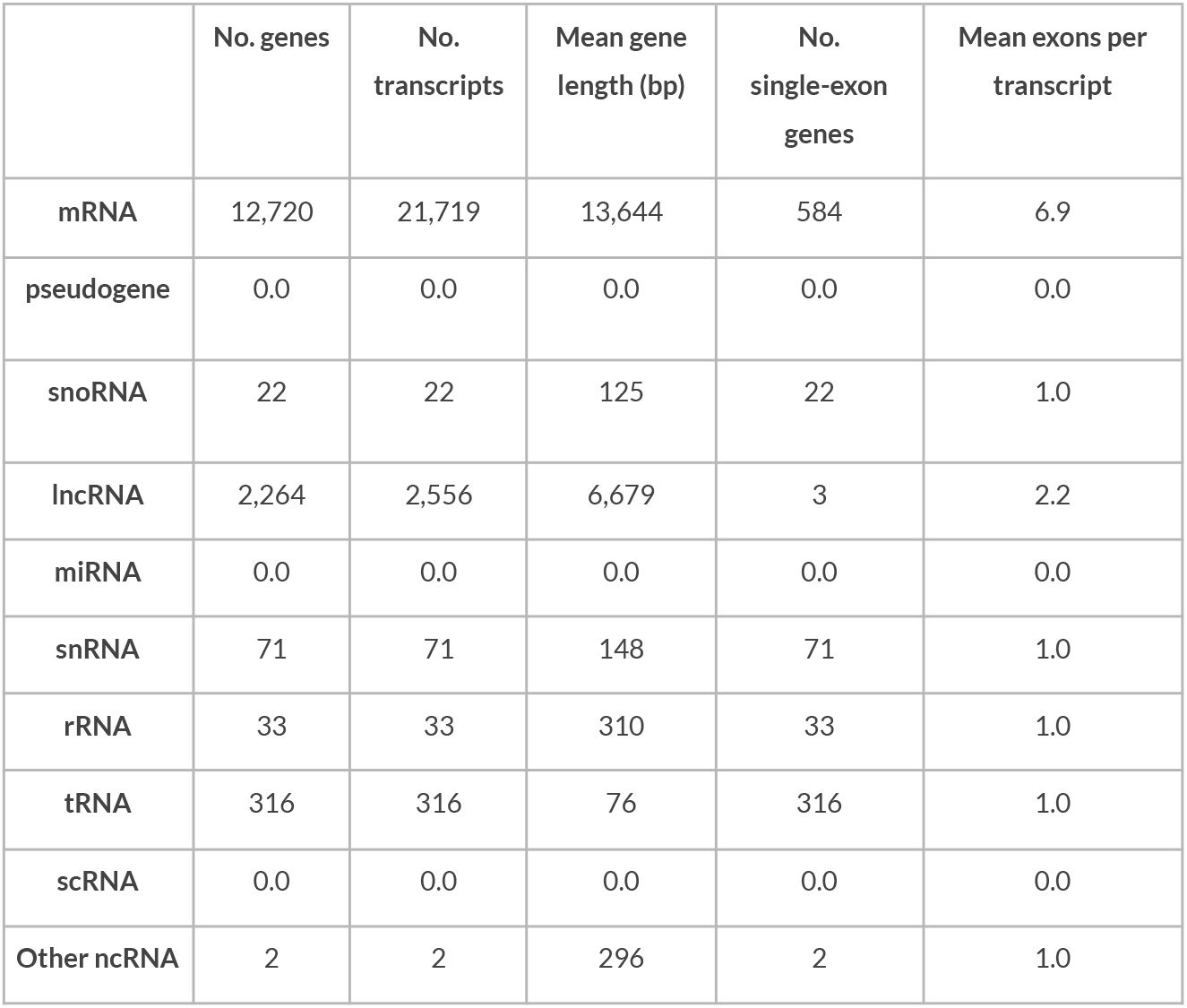
Statistics from assembled gene models.

**Table 2.** Annotation completeness and consistency scores calculated by BUSCO run in protein mode (lepidoptera_odb10) and OMArk (Metazoa-v2.0.0.h5)

|  | Complete | Single copy | Duplicated | Fragmented | Missing |
| --- | --- | --- | --- | --- | --- |
| BUSCO | 4,925 (93.2%) | 4,878 (92.3%) | 47 (0.9%) | 104 (2.0%) | 257 (4.8%) |
| OMArk | 7,301 (92.0%) | 6,844 (86.3%) | 457 (5.7%) | - | 627 (7.9%) |
|  | Consistent | Inconsistent | Contaminants | Unknown |  |
| OMArk | 11,434 (89.9%) | 363 (2.8%) | 0.0 (0.0%) | 923 (7.2%) |  |

## Acknowledgements

We acknowledge the support of the Freiburg Galaxy Team: Saim Momin and Björn Grüning, Bioinformatics, University of Freiburg (Germany), funded by the German Federal Ministry of Education and Research BMBF grant 031 A538A de.NBI-RBC and the Ministry of Science, Research and the Arts Baden-Württemberg (MWK) within the framework of LIBIS/de.NBI Freiburg. We would like to acknowledge the assembly reviewer, Emilie Teodori, from Genoscope.

## Author Contributions

TČ and EB coordinated the project; TČ collected the species; TČ identified the species; TČ sampled and preserved biological material and provided metadata; RF, NE, AsB, RM, and TL provided support in sampling, shipping of biological material, metadata collection, and management; LA and MG extracted DNA, prepared libraries and performed sequencing; FCF, FC, and JGG performed genome assembly and curation under the supervision of TA; LH, SS, and FM performed genome annotation; CB generated the analysis and report. All authors contributed to the writing, reviewing, and editing of this genome note and read and approved the final version.

## Literature Cited

Aken, B. L., Ayling, S., Barrell, D., Clarke, L., Curwen, V., Fairley, S., Fernandez Banet, J., Billis, K., García Girón, C., & Hourlier, T. (2016). The Ensembl gene annotation system. Database, 2016, baw093.

Article 17 Reporting, 2013–2018. Reporting under Article 17 of the Habitats Directive. https://nature-art17.eionet.europa.eu/article17/

Čelik, T., Bräu, M., Bonelli, S., Cerrato, C., Vreš, B., Balletto, E., Stettmer, C., & Dolek, M. (2015). Winter-green host-plants, litter quantity and vegetation structure are key determinants of habitat quality for Coenonympha oedippus in Europe. Journal of Insect Conservation, 19, 359–375.

Čelik, T., Kralj-Fišer, S., Šilc, U., Küzmič, F., & Vreš, B. (2024). Reviving of Coenonympha oedippus: A comprehensive approach to the reintroduction of an endangered European butterfly. Insect Conservation and Diversity, icad.12778.

Čelik, T., Küzmič, F., Šilc, U., & Vreš, B. (2021). Raziskava stanja potencialnih izvornih populacij vrste barjanski okarček (Coenonympha oedippus) in stanja njihovega habitata s smernicami za ustrezno upravljanje: Končno poročilo. Biološki inštitut Jovana HadŽija ZRC SAZU.

Challis, R., Kumar, S., Sotero-Caio, C., Brown, M., & Blaxter, M. (2023). Genomes on a Tree (GoaT): A versatile, scalable search engine for genomic and sequencing project metadata across the eukaryotic tree of life. Wellcome Open Research, 8, 24.

Consortium, U. (2019). UniProt: A worldwide hub of protein knowledge. Nucleic Acids Research, 47(D1), D506–D515.

Council Directive 92/43/EEC of 21 May 1992 on the conservation of natural habitats and of wild fauna and flora.

Després, L., Henniaux, C., Rioux, D., Capblancq, T., Zupan, S., Čelik, T., Sielezniew, M., Bonato, L., & Ficetola, G. F. (2019). Inferring the biogeography and demographic history of an endangered butterfly in Europe from multilocus markers. Biological Journal of the Linnean Society, 126(1), 95–113.

Dierks, K. (2006). Beobachtungen zur Larvalbiologie von Coenonympha oedippus (Fabricius, 1787) im Südwesten Frankreichs (Lepidoptera: Satyridae). Entomologische Zeitschrift, 116(4), 186–188.

Gomez-Garrido, J. (2024). CLAWS (CNAG’s Long-read Assembly Workflow in Snakemake) [Computer software].

Greenwood, M. P., Capblancq, T., Wahlberg, N., & Després, L. (2025). Whole genome data confirm pervasive gene discordance in the evolutionary history of Coenonympha (Nymphalidae) butterflies. Molecular Phylogenetics and Evolution, 202, 108222.

Gruber, A. R., Lorenz, R., Bernhart, S. H., Neuböck, R., & Hofacker, I. L. (2008). The Vienna RNA websuite. Nucleic Acids Research, 36(suppl_2), W70–W74.

Jugovic, J., Zupan, S., BuŽan, E., & Čelik, T. (2018). Variation in the morphology of the wings of the endangered grass-feeding butterfly Coenonympha oedippus (Lepidoptera: Nymphalidae) in response to contrasting habitats. European Journal of Entomology, 115.

Kalvari, I., Nawrocki, E. P., Argasinska, J., Quinones-Olvera, N., Finn, R. D., Bateman, A., & Petrov, A. I. (2018). Non-Coding RNA Analysis Using the Rfam Database. Current Protocols in Bioinformatics, 62(1), e51.

Kozomara, A., Birgaoanu, M., & Griffiths-Jones, S. (2019). miRBase: From microRNA sequences to function. Nucleic Acids Research, 47(D1), D155–D162.

Lhonoré, J. (1998). Biologie, écologie et répartition de quatre espèces de Lépidoptères Rhopalocères protégés (Lycaenidae, Satyridae) dans l’ouest de la France. OPIE.

Manni, M., Berkeley, M. R., Seppey, M., Simão, F. A., & Zdobnov, E. M. (2021). BUSCO update: Novel and streamlined workflows along with broader and deeper phylogenetic coverage for scoring of eukaryotic, prokaryotic, and viral genomes. Molecular Biology and Evolution, 38(10), 4647–4654.

Mazzoni, C. J., Ciofi, C., & Waterhouse, R. M. (2023). Biodiversity: An atlas of European reference genomes. Nature, 619, 252–252.

Nawrocki, E. P., & Eddy, S. R. (2013). Infernal 1.1: 100-fold faster RNA homology searches. Bioinformatics, 29(22), 2933–2935.

Nevers, Y., Warwick Vesztrocy, A., Rossier, V., Train, C.-M., Altenhoff, A., Dessimoz, C., & Glover, N. M. (2025). Quality assessment of gene repertoire annotations with OMArk. Nature Biotechnology, 43(1), 124–133.

Rhie, A., Walenz, B. P., Koren, S., & Phillippy, A. M. (2020). Merqury: Reference-free quality, completeness, and phasing assessment for genome assemblies. Genome Biology, 21(1), 245.

van Swaay, C. (2010). Coenonympha oedippus (Europe assessment). The IUCN Red List of Threatened Species.

Zupan, S., Jugovic, J., Čelik, T., & Buzan, E. (2021). Population genetic structure of the highly endangered butterfly Coenonympha oedippus (Nymphalidae: Satyrinae) at its southern edge of distribution. Genetica, 149(1), 21–36.

